# OncotreeVIS – an interactive graphical user interface for visualizing mutation tree cohorts

**DOI:** 10.1101/2024.11.15.623847

**Authors:** Monica-Andreea Baciu-Drăgan, Niko Beerenwinkel

## Abstract

**Motivation:** In recent years, developments in single-cell next-generation sequencing technology and computational methodology have made it possible to reconstruct, with increasing precision, the evolutionary history of tumors and their cell phylogenies, represented as mutation trees. Many mutation tree inference tools exist, but they do not support detailed visual tree inspection, nor tree comparisons or analysis at the cohort level, an important task in computational oncology.

**Results:** We developed oncotreeVIS, an interactive graphical user interface for visualizing mutation tree cohorts and tree posterior distributions obtained from mutation tree inference tools. OncotreeVIS can display mutation trees that encode single or joint genetic events, such as point mutations and copy number changes, and highlight matching subclones, conserved trajectories and drug-gene interactions at cohort level. OncotreeVIS facilitates the visual inspection of mutation tree clusters and pairwise tree distances. It is available both as a JavaScript library that can be used locally or as a web application that can be accessed online. It includes seven default datasets of public mutation tree cohorts for visualization, while new mutation trees are provided in a predefined JSON format.

**Availability and implementation:** https://github.com/cbg-ethz/oncotreeVIS.

**Web application:** https://cbg-ethz.github.io/oncotreeVIS.

## 1 Introduction

Tumor progression is a process driven by the accumulation of genomic mutations, such as point mutations or copy number alterations (CNA) and shaped by environmental selection pressures, including those exhibited by immune responses and drug treatments [28]. This somatic evolutionary process can lead to a high level of heterogeneity in tumors both in space and time. Furthermore, it has been shown that the temporal order in which mutations occur can have a significant impact on clinical patient outcome [22, 21].

However, tracking the evolution of tumors over time is impossible before the point when the patient is being diagnosed and usually also not possible for solid tumors, where samples are available only through biopsy, when the tumor is removed from the patient. In these cases when the evolution of the tumor cannot be directly observed, it can be inferred computationally from the genetic profiles detected at the time of the biopsy in different cell populations in the tumor (subclones), using mutation tree inference tools such as SCITE [11], SPRUCE [8], or CONIPHER [10] for point mutations from single-cell, bulk and multi-sample data, respectively, SCICoNE [16] for copy number events, or COMPASS for joint copy number and point mutations [26].

The mutation tree inference tools process the genomic sequencing data and reconstruct its phylogeny by a probabilistic approach using as taxa either the individual cells (for single-cell data) or the clones inferred based on variant allele frequencies, for bulk data [14].

The mutational history of a tumor can be represented by a mutation tree, also referred to as *event tree*, where the nodes correspond to different genetic events (event nodes) and the tree structure encodes the partial temporal order in which they occurred. The tumor cells are then attached to the mutation nodes, and their genetic profile is obtained by tracking all the events from the attachment node back to the root.

Depending on the statistical models used, their assumptions and their capability of handling errors in the real data, the results of different mutation tree inference tools do not always agree. They also depend on the resolution at which the input data is sequenced (bulk or single-cell), on the amount of cells per sample, and on the number of samples extracted from different sites of the biopsy.

Once reconstructed, the individual mutation trees are either visually inspected in order to understand the intra-tumor heterogeneity and potentially guide clinical decisions in tumor boards, or they can be analyzed in a comparative fashion as tree cohorts in order to compute conserved evolutionary trajectories [5, 17] or mutation tree embeddings and pairwise tree distance scores [2]. A visualization of the mutation trees is typically provided as part of the output of tree inference tools in Graphviz format [9]. More complex visualization tools for clonal evolution take into account longitudinal input data (i.e., data sampled at successive time points in the life of the tumor) and treatment information, and are able to generate fish plot views of the tumors [18, 25].

However, the existing mutation tree visualization options are generated locally, are limited to individual trees, and either display all or none of the subclonal genomic event annotations in the mutation tree nodes. This makes the visualization less informative when the annotations are discarded and overcrowded when the set of affected genes per subclone is large. Also, all existing tree visualization tools are provided as code libraries for different programming languages and need local installation and code integration in order to be used, which makes the process of generating and sharing results more difficult. Nonetheless, no tools for comparative tree visualization of cohorts exist.

To address these limitations, we developed oncotreeVIS, an interactive web-based graphical user interface for visualizing mutation trees from large cohorts, which helps assess heterogeneity among tumors. The trees can be displayed side by side, grouped according to a given clustering, or projected into a 2D latent space given the pairwise tree distances. Additional information such as clinical data, drug-gene interactions, matching mutation patterns and conserved trajectories between groups of trees, and k-nearest neighbor trees, is displayed in an interactive fashion.

OncotreeVIS can be effectively used to visually inspect the results of mutation tree clustering methods, the tree posterior distributions of mutation tree inference tools, or different versions of mutation trees for the same patient, e.g., mutation trees reconstructed with different mutation tree inference tools, or longitudinal samples.

OncotreeVIS is available both as a JavaScript library and as an online web application, which makes it easy to use both for computational scientists and clinicians. By default, oncotreeVIS provides visualizations of seven publicly available mutation tree cohorts reconstructed with existing mutation tree inference tools from both bulk and single-cell data which highlight some of the use cases: visualizing matching patterns within clustered point mutation and CN trees for different cancer types [19, 26, 27, 23, 13], visualizing tree structures corresponding to different modes of spatial tumor evolution (linear, branching, punctuated, [20]), inspecting conserved trajectories among alternative trees sampled from the posterior tree distribution [1] (see Table 1 and Suppl. Fig. 4 for details). New tree cohort datasets can be visualized easily by providing the data in a predefined JSON format.

**Table 1:** Public mutation tree cohorts visualized by default in the oncotreeVIS web application. The details of the tree cohort from Noble et al. 2022 are not reported because it is a mixture of 5 datasets. Abbreviations: single-cell (SC), multi-region (MR), whole-genome sequencing (WGS), whole exome sequencing (WES).

| Dataset | Description | Sequencing type | Mutation tree inference tool |
| --- | --- | --- | --- |
| Morita et al. 2020 [19] | 153 AML point mutation trees from 123 patients | SC, panel | SCITE [12] |
| Sollier et al. 2023 [26] | 145 AML joint CN and point mutation trees (same raw data as [19]) | SC, panel | COMPASS [26] |
| Wegmann et al. 2024 [27] | 25 CN trees from 16 patients | SC, WGS | SCICoNE [16] |
| Razavi et al. 2018 [23] | 1,214 breast cancer mutation trees (1,152 patients) preprocessed following [6] | Bulk | SPRUCE [8] |
| TRACERx [13] | 99 point mutation trees from non-small cell lung cancer patients | MS, WES | Jamal-Hanjani et al. 2017 [13] |
| TRACERx421 [1] | 4,843 alternative phylogenies inferred for 126 non-small cell lung metastatic cancer patients | MS, WES | CONIPHER [10] |
| Noble et al. 2022 [20] | 43 mutation trees with different modes of evolution, from 5 cancer types | NA | NA |

## 2 Usage and implementation details

OncotreeVIS is implemented in HTML5 and JavaScript, and uses the D3.js library for data visualization [3]. To render large amounts of trees without blocking the browser it performs on the fly computations asynchronously and uses lazy loading of the HTML elements that are offscreen.

The input mutation trees are provided as a JSON object which encodes the tree structures with node labels representing single or joint genomic events acquired by each subclone at different times in the inferred evolution of the tumor. The mutation events in each node are encoded by a dictionary where the keys are the type of event (point mutation, CNA, etc.) and the value stores additional information, e.g., the point mutation variant, or the copy number amplification/deletion amounts w.r.t. the diploid state. Additional node annotations can be provided by the user: matching label IDs, which uniquely identify the set of mutation events and are used to facilitate the identification of matching subclones, subclone sizes, and clinical information if available, as described in Suppl. Table 4. The node sizes reflect the relative size of the subclonal cell populations. To improve the user experience we chose to use equal branch lengths for all the trees and display the number of mutations and their type as branch labels.

In our code repository we provide scripts to convert the output formats of four common mutation tree inference tools (Suppl. Table 4) into the JSON input format used by oncotreeVIS. In addition to individual mutation trees, oncotreeVIS can incorporate a given tree clustering and a precomputed pairwise tree distance matrix, which can be provided as additional key value pairs in the JSON input file. For the predefined tree cohorts, we used oncotree2vec [2] to compute the pairwise distances and the clustering of the mutation trees based on the matching pairwise relations between the clones.

## 3 Features

In order to explore the heterogeneity of mutation tree cohorts, oncotreeVIS provides an interactive interface where the user can navigate between different views of the data both at the tree and cohort level. In addition, one can visualize subclones with matching genetic events, conserved evolutionary trajectories, and the impact of subclonal target gene events on drug efficacy (Suppl. Table 4).

### 3.1 Tree cohort views

OncotreeVIS provides different ways of visualizing event trees from large mutation tree cohorts. For each tree, we provide text labels summarizing the mutation events or (for more than two events) the number of mutation events of each type acquired in each subclone node (Fig. 1A). By default, the individual trees are displayed side by side. If a clustering is provided, the trees are ordered according to the given cluster groups, which are indicated by different background colors (Fig. 1B). The user can choose whether to highlight matching subclones (i.e., the nodes with the same matching label ID) or trajectories (i.e., tree branches) conserved between the trees from the same cluster. If no clustering is provided, the matching subclones across the entire cohort will be highlighted by node colors. Detailed information is displayed by clicking on the individual mutation trees (see Section 3.2 below).

**Figure 1:**
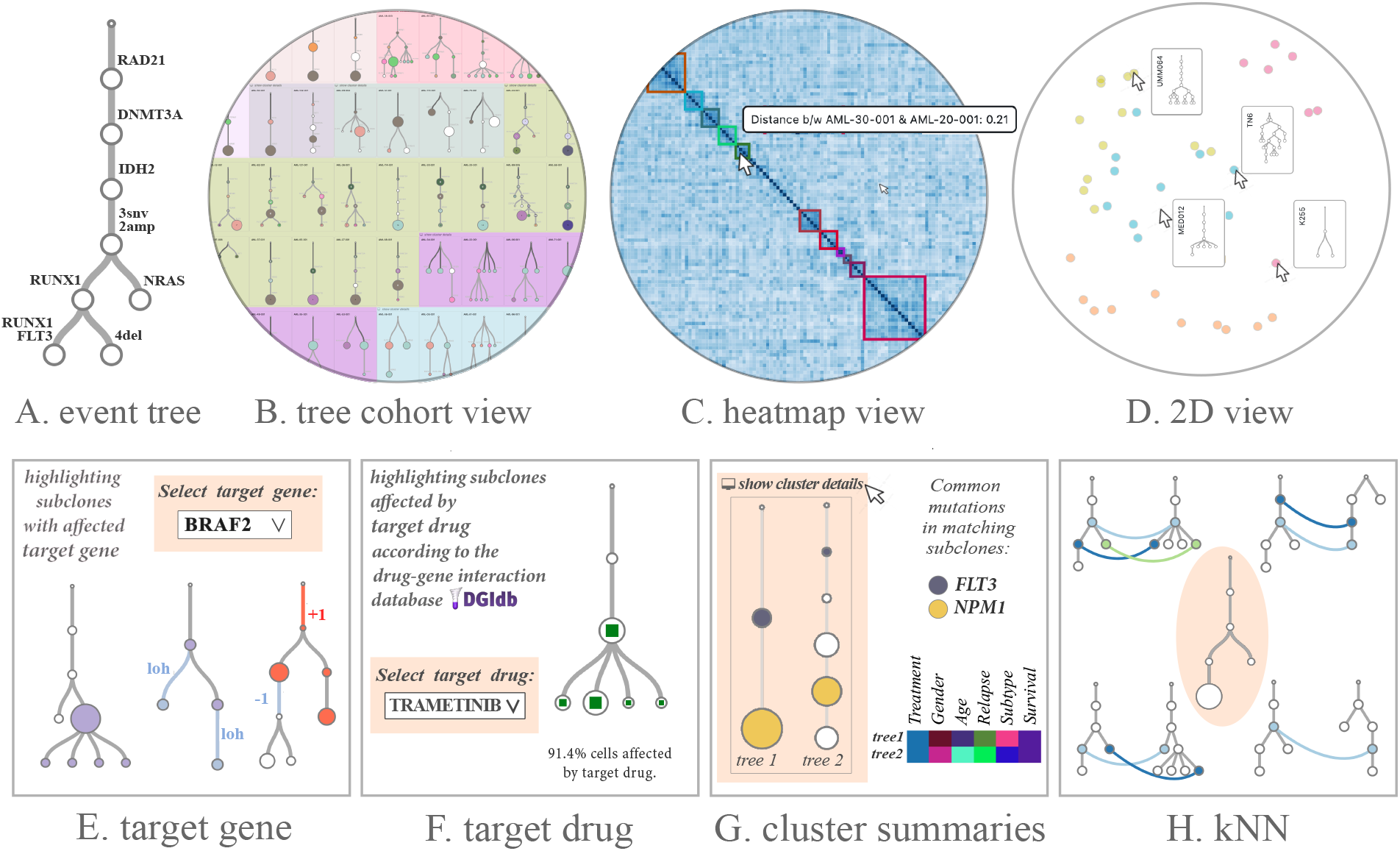
Screenshots of different mutation tree views provided by oncotreeVIS. Some interactive elements are indicated by the arrow pointer. **(A)** Mutation tree with events acquired in each subclone node labeled on the incoming edge (e.g., “3snv, 2amp, 4del” stands for three point mutations, two copy number amplifications, and four deletion w.r.t. the diploid state). **B–D:** Three different tree cohort views: **(B)** Mutation trees displayed side by side, grouped by a given clustering; matching subclones and conserved trajectories are highlighted. **(C)** Heatmap visualization of the pairwise distances between the mutation trees. **(D)** 2D visualization of the mutation trees based on the pairwise distances, using Multidimensional scaling (MDS); each tree is a dot in the latent space and can be visualized when the dot is hovered by the mouse pointer. E–H: Additional information at mutation tree level: **(E)** Highlighting subclones with an affected target gene using red color for CN amplification, blue for CN deletion and violet for any other mutation event. **(F)** The subclones affected by a selected target drug are indicated by the green squares; there is one clone (next to the root) which does not interact with the selected drug. **(G)** Cluster summaries (clinical data and matching subclonal events). **(H)** K-nearest neighbors of a target mutation tree (highlighted with orange background) among other trees in the cohort.

Two additional cohort views are available if the pairwise distances between the mutation trees are provided by the user: a heatmap view of the tree distances and a 2D representation of the trees in latent space, computed on the fly using Multidimensional scaling (Fig. 1C,D). Similarly to the tree view, the different tree clusters are highlighted if a clustering is provided.

By clicking on the clusters, additional details are displayed: a summary of the clinical data and information on the mutation events shared between the subclones with the same matching label ID in the cluster trees (Fig. 1G).

### 3.2 Mutation tree information

In order to facilitate the visual inspection of the mutation trees for guiding clinical decisions, oncotreeVIS provides an interactive user interface where the user can track the impact of certain genes and drugs of interest on the evolution of the tumors (Fig. 1E,F). For each selected tree, the user can choose from the list of genes affected by mutation events in the tree to visualize the subclone (i.e., tree node) which first acquired that mutation and consult the list of drugs which interact with the target gene according to the Drug Gene Interaction database (DGIdb, [4]), where only drug-gene interactions with more than three citations are considered. Some of these mutated genes can be targeted by specific drugs and one can select a drug and visualize the subclones which contain affected genes which interact with the target drug according to DGIdb. For this task, we compute the mutation profile of each subclone by collecting all mutation events along the path from the root to any node in the mutation tree.

Tracking the mutation events which affect a specific target gene in the mutation tree can provide insights into its role in tumor growth and therapeutic resistance. Also, by selecting a target drug the user can make sense of the fraction of the tumor that is potentially impacted by the drug and can help find new treatment strategies.

### 3.3 K-nearest tree neighbors

A key question in cancer genomics is to identify recurring patterns of tumor evolution. Mutation tree cohorts can reveal such patterns of the order in which mutations accumulate in different trees, especially co-occurring and mutually exclusive mutation events, which have an essential role in shaping tumor progression [15]. The problem of finding such patterns has been addressed by several existing tree distance metrics which focus on different tree features such as common ancestor or distinctly inherited sets [7], number of edge-induced partitions [12], or pairwise node relations [2]. OncotreeVIS incorporates the information about the pairwise distances between the mutation trees, if provided by the user, to project the trees in a 2D latent space. In the absence of a precomputed tree distance matrix we propose a greedy heuristic (Suppl. Algorithm 1, Suppl. Figure 4) that approximates the problem of maximum matching with ordering constraints, which is NP-hard [24], and computes on the fly the nearest neighbors based on the provided matching labels. The matching trees are then displayed as an extension of the mutation tree information box.

## 4 Conclusions

OncotreeVIS is an interactive user interface to visualize mutation tree cohorts in order to assess the heterogeneity within and among tumors, the tree posterior distributions computed by different tree inference tools, pairwise tree distances, predicted drug-tumor interactions, and different modes of evolution. Our visualization can support clinical decision making and enhance research on the genetic basis of tumor progression. OncotreeVIS is freely available under MIT license and can be used locally and integrated into more complex pipelines using the JavaScript library. Furthermore, we provide an online web application where everyone with a browser can visualize seven predefined public mutation tree cohorts or load their own data on the fly.

## Acknowledgements

The authors would like to thank my peers from the Computational Biology Group at ETH Zurich (Diane Duroux, Kerstin Lenhof and Johannes Gawron) for testing and providing feedback on oncotreeVIS and Caitlin Harrigan for making us aware of the CONIPHER mutation tree dataset.

## Supplementary information

**Supplementary Table 1.**
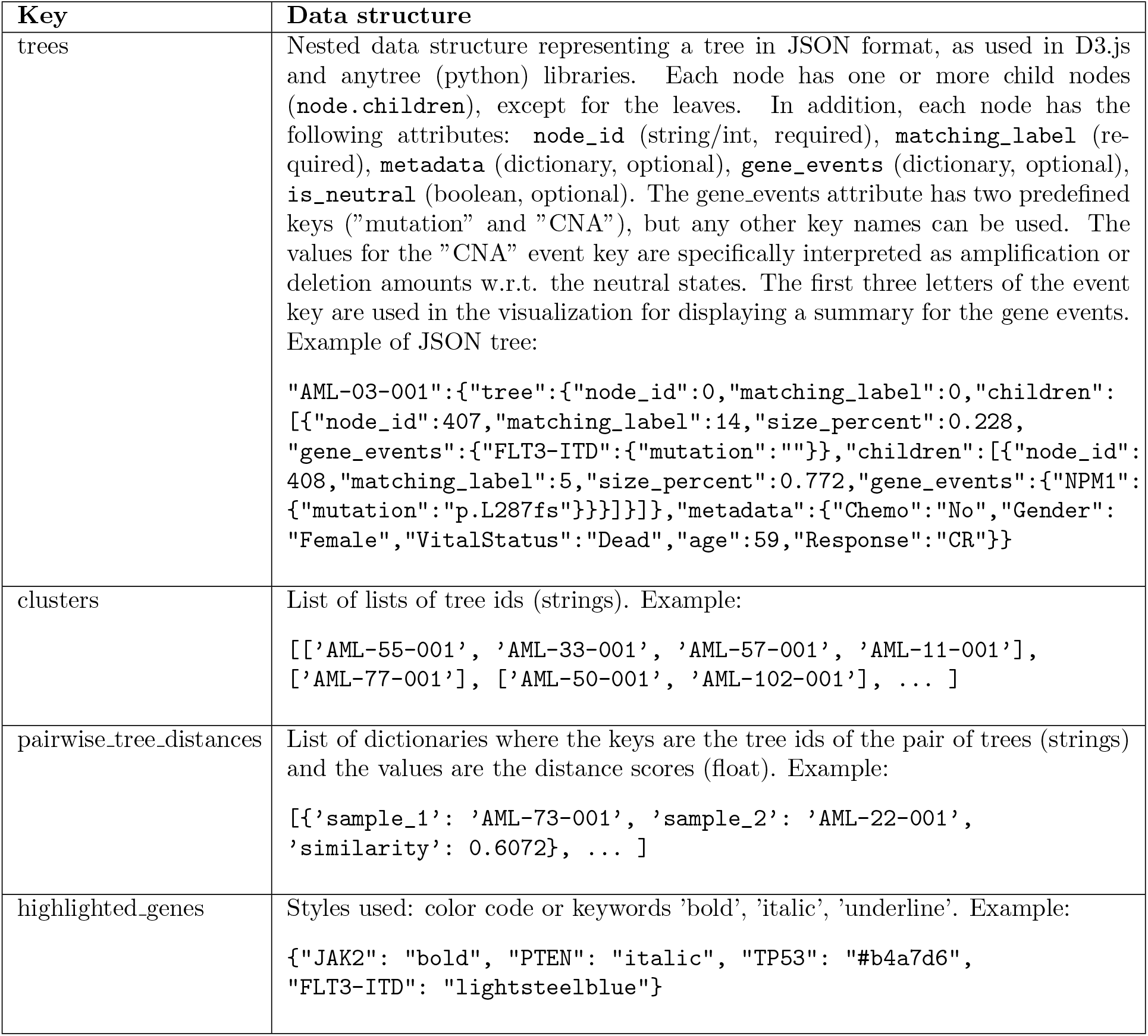
Example of input data encoding the information about a tree cohort in JSON format. Key values are highlighted in bold. Scripts for converting the output of different tree inference tools (see Suppl. Table 4) into the JSON format used by oncotreeVIS JSON are provided at https://github.com/cbg-ethz/oncotreevis.

**Supplementary Table 2.**
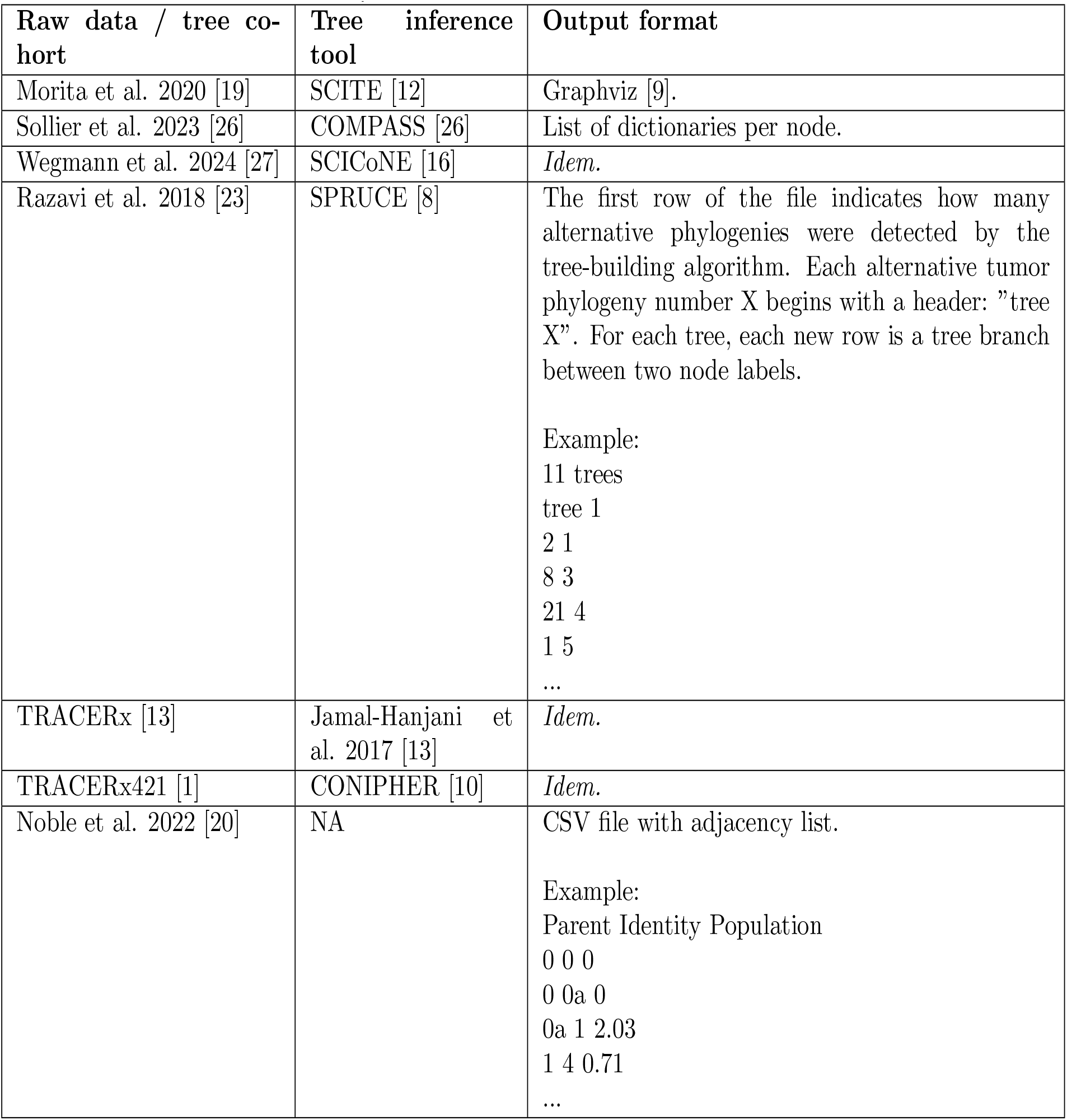
Output formats for the mutation trees output by different tree inference tools referred to in the paper. In our code repository we provide scripts to convert all these data formats into the JSON input format used by oncotreeVIS.

| Raw data / tree cohort | Tree inference tool | Output format |
| --- | --- | --- |
| Morita et al. 2020 [19] | SCITE [12] | Graphviz [9]. |
| Sollier et al. 2023 [26] | COMPASS [26] | List of dictionaries per node. |
| Wegmann et al. 2024 [27] | SCICoNE [16] | *Idem.* |
| Razavi et al. 2018 [23] | SPRUCE [8] | The first row of the file indicates how many alternative phylogenies were detected by the tree-building algorithm. Each alternative tumor phylogeny number X begins with a header: "tree X". For each tree, each new row is a tree branch between two node labels.<br>Example:<br>11 trees<br>tree 1<br>2 1<br>8 3<br>21 4<br>1 5<br>... |
| TRACERx [13] | Jamal-Hanjani et al. 2017 [13] | *Idem.* |
| TRACERx421 [1] | CONIPHER [10] | *Idem.* |
| Noble et al. 2022 [20] | NA | CSV file with adjacency list.<br>Example:<br>Parent Identity Population<br>0 0 0<br>0 0a 0<br>0a 1 2.03<br>1 4 0.71<br>... |

**Supplementary Table 3.**
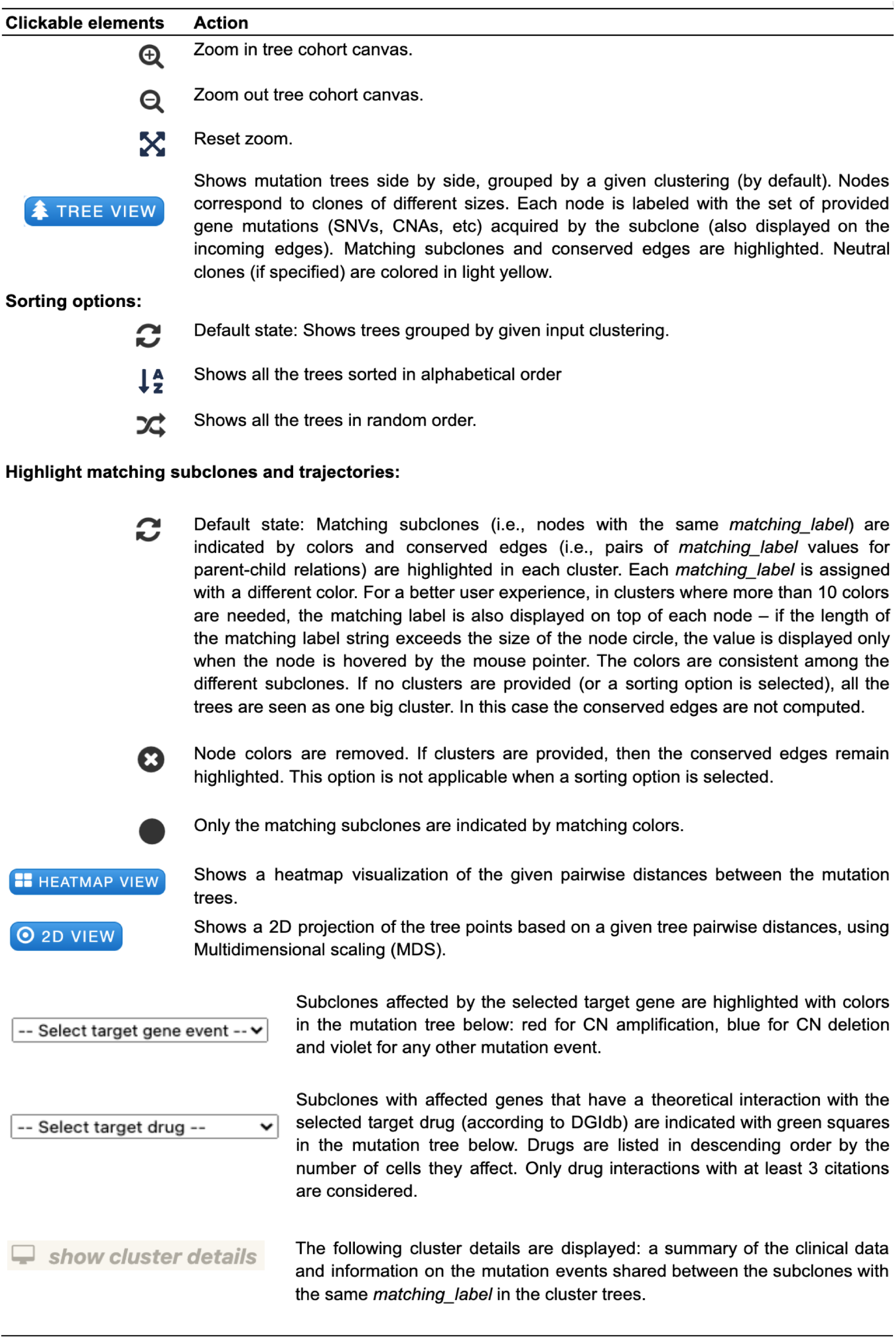
Implementation details.

| Clickable elements | Action |
| --- | --- |
|  | Zoom in tree cohort canvas. |
|  | Zoom out tree cohort canvas. |
|  | Reset zoom. |
| TREE VIEW | Shows mutation trees side by side, grouped by a given clustering (by default). Nodes correspond to clones of different sizes. Each node is labeled with the set of provided gene mutations (SNVs, CNAs, etc) acquired by the subclone (also displayed on the incoming edges). Matching subclones and conserved edges are highlighted. Neutral clones (if specified) are colored in light yellow. |
| **Sorting options:** |  |
|  | Default state: Shows trees grouped by given input clustering. |
|  | Shows all the trees sorted in alphabetical order |
|  | Shows all the trees in random order. |
| **Highlight matching subclones and trajectories:** |  |
|  | Default state: Matching subclones (i.e., nodes with the same *matching_label*) are indicated by colors and conserved edges (i.e., pairs of *matching_label* values for parent-child relations) are highlighted in each cluster. Each *matching_label* is assigned with a different color. For a better user experience, in clusters where more than 10 colors are needed, the matching label is also displayed on top of each node – if the length of the matching label string exceeds the size of the node circle, the value is displayed only when the node is hovered by the mouse pointer. The colors are consistent among the different subclones. If no clusters are provided (or a sorting option is selected), all the trees are seen as one big cluster. In this case the conserved edges are not computed. |
|  | Node colors are removed. If clusters are provided, then the conserved edges remain highlighted. This option is not applicable when a sorting option is selected. |
|  | Only the matching subclones are indicated by matching colors. |
| HEATMAP VIEW | Shows a heatmap visualization of the given pairwise distances between the mutation trees. |
| 2D VIEW | Shows a 2D projection of the tree points based on a given tree pairwise distances, using Multidimensional scaling (MDS). |
| -- Select target gene event -- | Subclones affected by the selected target gene are highlighted with colors in the mutation tree below: red for CN amplification, blue for CN deletion and violet for any other mutation event. |
| -- Select target drug -- | Subclones with affected genes that have a theoretical interaction with the selected target drug (according to DGIdb) are indicated with green squares in the mutation tree below. Drugs are listed in descending order by the number of cells they affect. Only drug interactions with at least 3 citations are considered. |
| show cluster details | The following cluster details are displayed: a summary of the clinical data and information on the mutation events shared between the subclones with the same *matching_label* in the cluster trees. |

**Supplementary Figure 1.**
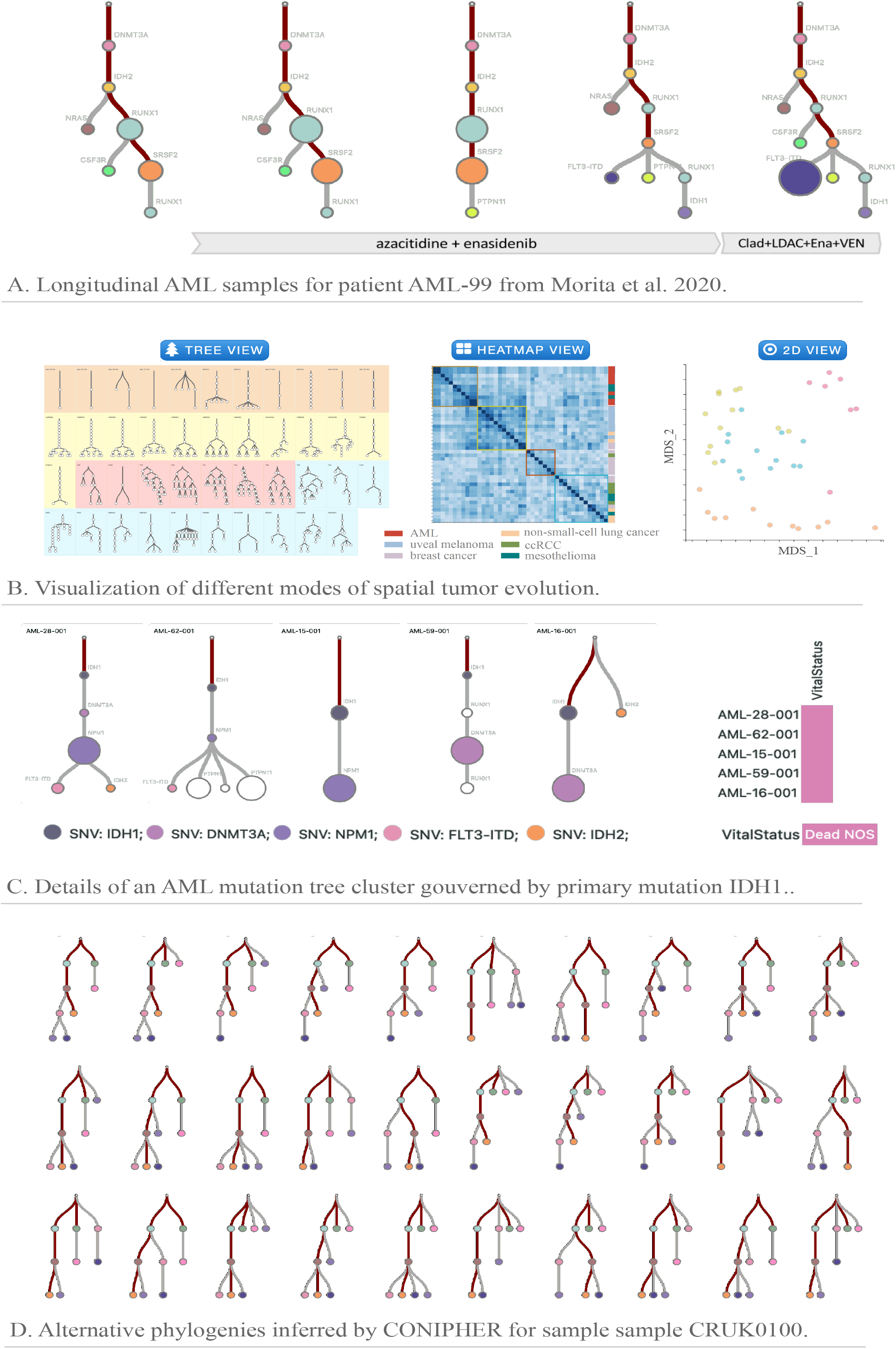
OncotreeVIS data views for four use cases: **(A)** Five longitudinal AML samples for patient AML-99 from Morita et al. 2020 obtained before and during treatment, including treatment change. The tumor evolution shown at different timepoints reveals the underlying process of therapeutic resistance, namely the emergence of IDH1/FLT3/NRAS clones during IDH2 inhibitor-containing therapy; **(B)** Cohort overview, heatmap, and 2D visualization of 43 tumor mutation trees from 6 cancer types and different modes of spatial tumor evolution selected in Noble et al. 2022, clustered with oncotree2vec (Baciu-Drăgan and Beerenwinkel 2024). The clusters are indicated by different colors and correspond to four modes of evolution: linear (orange cluster), punctuated (yellow cluster), branching (red cluster) and linear-to-branched evolution (blue cluster). **(C)** Details of an AML mutation tree cluster output by oncotree2vec, governed by primary mutation IDH2. **(D)** Alternative phylogenies inferred by CONIPHER mutation tree inference algorithm (Grigoriadis et al. 2024) for sample CRUK0100. Conserved trajectories between the alternative trees are highlighted in dark red.

**Supplementary Figure 2.**
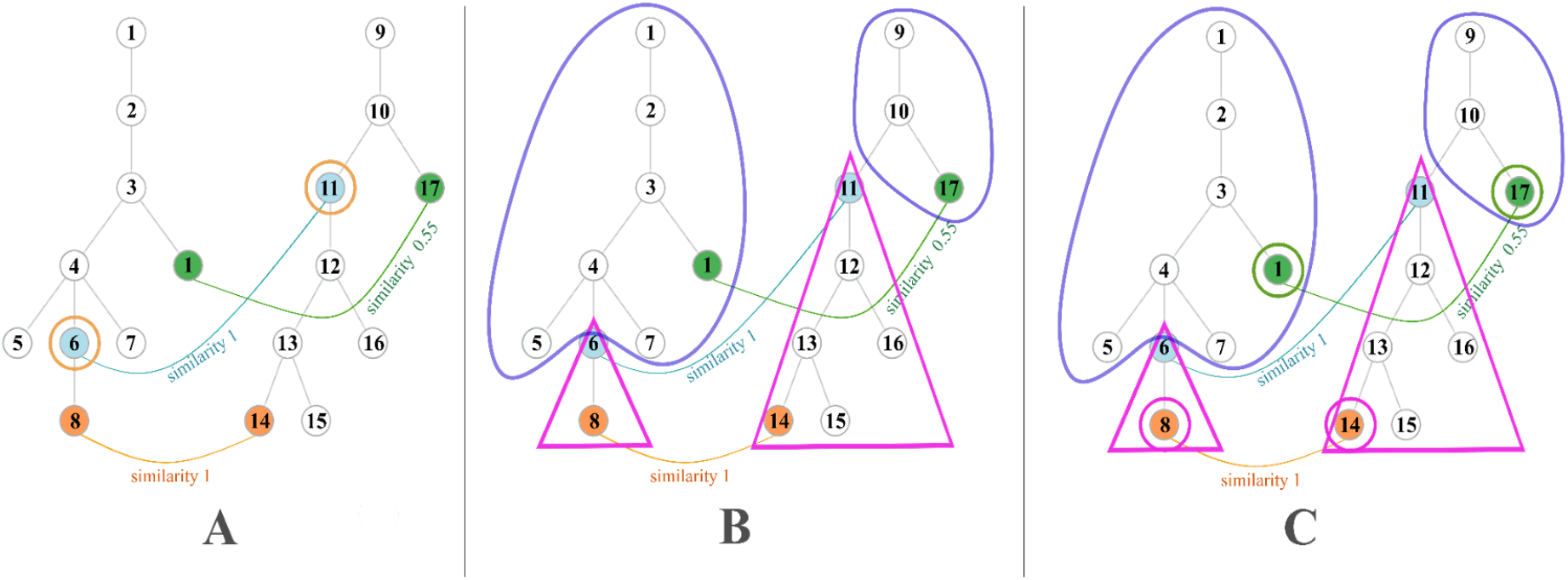
Example of how the proposed heuristic for the maximum matching problem with ordering constraints works for two trees with three matching nodes with similarities 1, 1 and 0.55. Each node has an unique id to make it easier to be referred. The nodes sharing the same color have matching label sets with a high similarity (the similarity score is specified along the corresponding link). White colored nodes are not matching any other node. **(A)** First, nodes with ids 6 and 11 are matched (maximum similarity and lowest depth). **(B)** The nodes from the child trees (the pink triangles) and from the outer trees (the blue triangles) are matched recurrently. **(C)** Inside the recursion, nodes with ids 8 and 14, and 1 and 17 respectively, are matched.

### Algorithm 1

A heuristic for the maximum matching with ordering constraints.

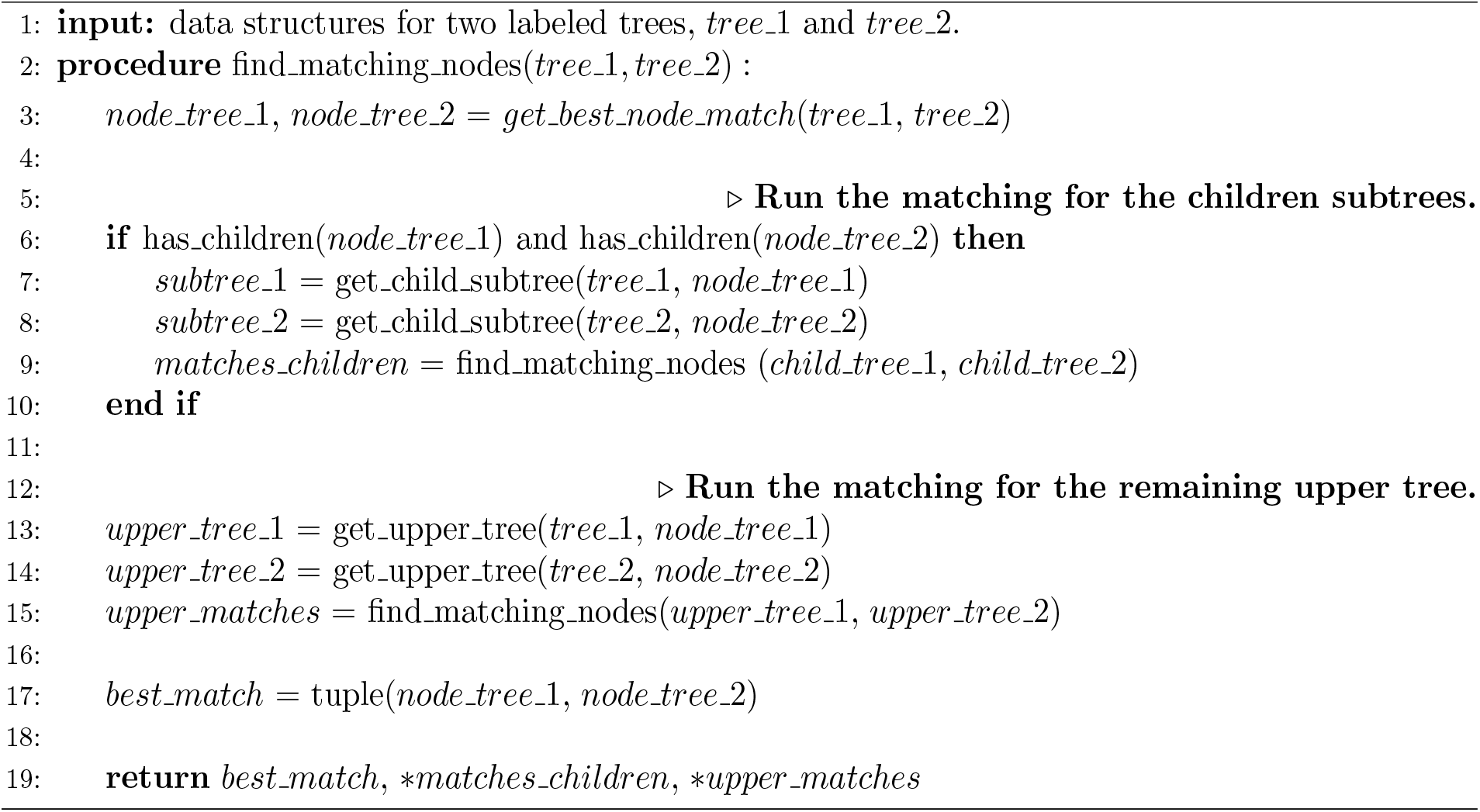

